# Cocaine-induced loss of LTD and social impairments are restored by fatty acid amide hydrolase inhibition

**DOI:** 10.1101/2023.06.05.543789

**Authors:** Laia Alegre-Zurano, Alba Caceres-Rodriguez, Paula Berbegal-Sáez, Olivier Lassalle, Olivier Manzoni, Olga Valverde

## Abstract

**Background:** A single usage of a drug of abuse can have lasting effects on both the brain and behavior, continuing even after the drug has been metabolized and eliminated from the body. A single dose of cocaine can abolish endocannabinoid-mediated long-term depression (eCB-LTD) in the nucleus accumbens (NAc) within 24 hours of administration. However, it is uncertain whether this altered neuroplasticity entails a behavioral deficit.

**Methods:** Our study employed adult male mice to investigate the effects of a single dose of cocaine (20 mg/kg) on eCB-LTD, saccharin preference, and social interactions 24 hours after administration. We also examined the gene expression in components of the eCB system. The pharmacological increase of anandamide was evaluated using the fatty acid amide hydrolase (FAAH) inhibitor URB597 (1 mg/kg).

**Results:** After a single dose of cocaine, mice displayed altered plasticity, social interactions, and preference for saccharin and a reduction in mRNA levels of the anandamide-catabolizing enzyme NAPE-PLD. We discovered that the FAAH inhibitor URB597 (1 mg/kg) successfully reversed the cocaine-induced loss of eCB-LTD in the NAc and restored normal social interaction in cocaine-exposed mice, but it did not affect their saccharin preference.

**Conclusions:** Overall, this research underlines the neuroplastic changes and subsequent behavioral alterations that occur after the initial use of cocaine, while also suggesting a potential role for anandamide in the early impairments caused by cocaine. The findings highlight the importance of understanding the mechanisms underlying the initiation of drug use and offer a potential therapeutic target.

## INTRODUCTION

The initiation of drug addiction stems from the first exposure to a rewarding substance. As drug consumption becomes frequent, the initial voluntary and goal-directed use can evolve into habitual and compulsive patterns, ultimately resulting in addiction in vulnerable people (1). Researchers have long studied the factors contributing to this transition (1), but even a single drug encounter can induce enduring changes in the central nervous system, persisting after the drug is no longer present (2,3).

The nucleus accumbens (NAc) plays a crucial role in drug-related and natural reward learning, with the endogenous cannabinoid (eCB) system (ECS) and synaptic plasticity in dopamine receptor 1 (D1)- and 2 (D2)-expressing medium spiny neurons (SPNs) implicated in reward-seeking behavior (4,5). The ECS comprises cannabinoid receptors 1 (CB1R) and 2 (CB2R), eCB ligands (anandamide, AEA, and 2-arachidonoylglycerol, 2-AG), and enzymes for synthesis (N-acylphosphatidylethanolamine-specific phospholipase D, NAPE-PLD, and diacylglycerol lipase, DAGL) and degradation (fatty acid amide hydrolase, FAAH, and monoacylglycerol lipase, MAGL) (6). In the NAc, the ECS modulates neuroplasticity in synapses through eCB-mediated long-term depression (eCB-LTD), a mechanism that downregulates excessive synaptic activity (7). eCB-LTD governs the strength of synapses from distal glutamatergic projections (e.g., the medial prefrontal cortex and the amygdala) onto SPNs (8). Activation of mGluR5 mediates eCB-LTD by triggering the synthesis of eCBs, which in turn induce LTD via two mechanisms: presynaptic inhibition of neurotransmitter release through CB1R activation (7), and postsynaptic internalization of α-amino-3-hydroxy-5-methyl-4-isoxazolepropionic acid receptors (AMPARs) through transient receptor potential vanilloid 1 (TRPV1) activation (8). While the specific eCB responsible for the cocaine-induced loss of plasticity is a subject of ongoing debate, AEA has emerged as a promising candidate because it activates CB1R (6) and possesses ideal characteristics for facilitating TRPV1-mediated plasticity, including its function as an intracellular messenger (9) and its full agonist activity at TRPV1 channels (10).

The molecular and physiological underpinnings of reward-seeking behavior remain incompletely understood. Pharmacological and genetic approaches have demonstrated that eCB-LTD in D1-expressing SPNs of the NAc is crucial for the expression of cocaine, natural reward (i.e., sucrose), and brain-stimulation-seeking behaviors (11). Rodents display various behavioral changes following cocaine exposure. However, it is unclear whether a single cocaine administration can induce behavioral changes beyond its acute effects. For example, repeated pairing of saccharin with cocaine decreases saccharin consumption (12,13). Similarly, both acute (14–16) and chronic (17) cocaine use in rodents reduces social interaction during drug use and even after one day of withdrawal from repeated exposure (18). However, although a single cocaine administration abolishes eCB-LTD in mice (3), it remains unclear how a single cocaine exposure affects NAc-related behaviors once the drug’s acute effects have subsided.

In this study, we report that a single dose of cocaine can alter behavior 24 hours after exposure, with disrupted social interactions and diminished saccharin preference. Additionally, our data suggest that elevating AEA levels through the FAAH inhibitor URB597 elicits beneficial effects, rescuing physiological eCB-LTD and adapted behavior. Therefore, we propose that the FAAH inhibitor URB597 could have a potential therapeutic effect for addressing initial impairments caused by cocaine.

## MATERIALS AND METHODS

### Animals

#### Behavior and biochemistry

Male C57BL/6 mice (postnatal day, PD, 56) were purchased from Janvier (Barcelona, Spain) and maintained in the animal facility (UBIOMEX, PRBB) in a 12-hour light-dark cycle. Mice were housed at a stable temperature (22 °C ± 2) and humidity (55% ± 10%), with food and water *ad libitum*. They were allowed to acclimatize to the new environmental conditions for at least five days prior to experimentation. Animal care and experimental protocols were approved by the Animal Ethics Committee (CEEA-PRBB),

#### Electrophysiology

Male C57Bl/6J mice (PD60) were purchased from Janvier (Le Genest-Saint-Isle, France) and maintained in the animal facility in a 12-hour light-dark cycle. Mice were housed at stable temperature (20 ± 1°C) and humidity (60%), with food and water *ad libitum*. They were allowed to acclimatize to the new environmental conditions for at least one week prior to experimentation. All experiments were performed on male C57BL/6J mice between PD70 and PD110. The French Ethical committee authorized this project (APAFIS#3279-2015121715284829 v5).

In all the studies, animals were treated in compliance with the European Communities Council Directive (86/609/EEC) for the care and use of laboratory animals.

### Materials

Cocaine HCl (20 mg/kg i.p.; Cocaine was kindly provided by the National Institute on Drug Abuse, Bethesda, MD; USA or purchased from Alcaliber S.A., Madrid, Spain) was dissolved in 0.9% NaCl. The FAAH inhibitor URB597 (1 mg/kg i.p. or 2µM) was purchased in Merck Life Science (Madrid, Spain) or Tocris Bioscience (Illkirch, France) and was dissolved in a vehicle solution composed by ethanol, cremophor E.L. (Sigma-Aldrich) and 0.9% NaCl (1:1:18).

### Experimental design

We used three different cohorts of mice (Fig. 1A and 1B). One cohort was used for the measurement of eCB-LTD. Mice were injected with saline or cocaine (20mg/kg) 24 hours before being euthanized and extracellular recordings of the NAc core were measured with the presence or absence of URB597. A second cohort of mice was used to test the effects of cocaine and URB597 on saccharin preference. After the habituation phase, mice were injected with either cocaine or saline, and 24 hours later with URB597 or vehicle. A third cohort of mice was used to test the effects of cocaine and URB597 on social behavior in the three-chamber social interaction test and to assess the mRNA levels of different components that participate in the eCB-LTD. Mice were pre-treated with cocaine or saline 24 hours before the test, and with URB597 or vehicle 30 minutes before the test. Immediately after the test, mice were euthanized, and the ventral striatum (vSTR) was dissected for qPCR analysis.

**Figure 1.**
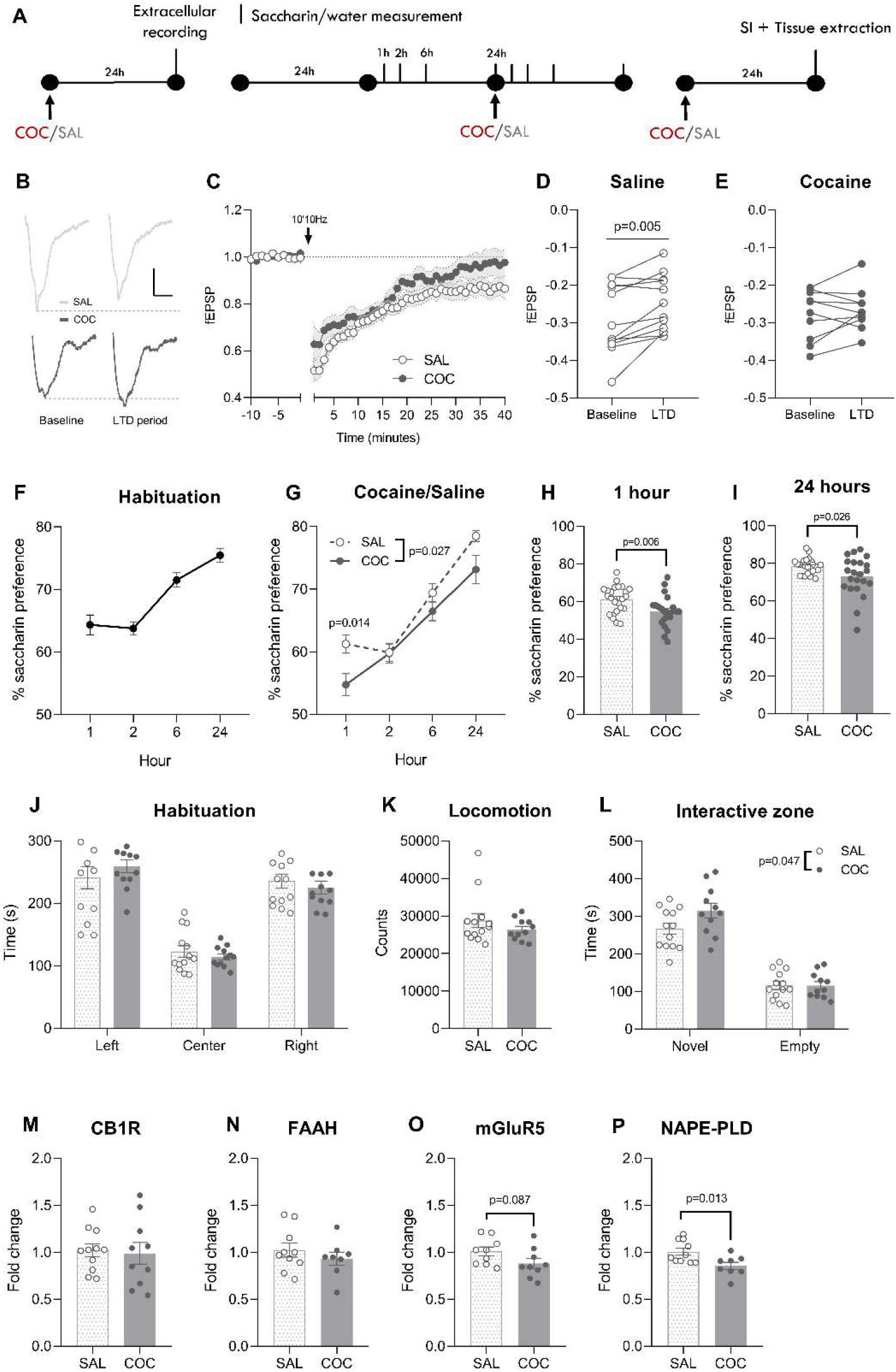
Single cocaine (20 mg/kg) exposure perturbates NAc core plasticity, related behaviors and NAPE-PDL gene expression. **A** Mice were treated with cocaine/saline and 24 hours later were tested for eCB-LTD in the NAc, saccharin preference, and social behavior and gene expression levels. **B** Representative traces during baseline (−10 to 0 minutes) and LTD expression period (30 to 40 minutes). Scale: 0.1 mV, 5 ms. **C** Average time courses of eCB-LTD 24 hours after receiving either saline (n=12) or cocaine (n= 10). **D** fECSP during baseline and LTD expression period in saline and **E** cocaine groups. **F** Preference for saccharin measured for 24 hours following cocaine/saline administration. During the habituation day and the test day, the measurements were performed at 1, 2, 6 and 24 hours. **G** Cocaine administration decreased preference for saccharin during the following day, especially after **H** 1 and **I** 24 hours (SAL n=24, COC n=22). **J** Time spent in each compartment during the habituation phase in the three-chamber social interaction test. Mice did show preference for the left/right compartment (SAL n=13, COC n=12). **K** Cocaine-treated mice did not differ in their locomotor activity during the habituation phase. **L** Time spent in the interactive zones surrounding the novel mouse or the empty holder during the sociability phase. Cocaine-treated mice showed an increase in social interaction. **M** Gene expression levels of CB1R, **N** FAAH, **O** mGluR5 and **P** NAPE-PLD in the vSTR of saline- (n=9-11) and cocaine-treated (n=8-10) after the social interaction test. SAL: saline, COC: cocaine, SI: social interaction, LTD: long term depression, fEPSP: field postsynaptic excitatory potential, CB1R: cannabinoid receptor 1, FAAH: fatty acid amide hydrolase, mGluR5: metabotropic glutamate receptor 5, NAPE-PLD: N-acylphosphatidylethanolamine-specific phospholipase D.

### Slice preparation and electrophysiology

Adult male mice were deeply anesthetized with isoflurane and sacrificed as previously described (7,19–21). The brain was sliced (300 μm) on the coronal plane with a vibratome (Integraslice, Campden Instruments) in a sucrose-based solution at 4°C (in mm as follows: 87 NaCl, 75 sucrose, 25 glucose, 2.5 KCl, 4 MgCl_2_, 0.5 CaCl_2_, 23 NaHCO_3_ and 1.25 NaH_2_PO_4_). Immediately after cutting, slices containing the NAc were stored for 1 h at 32°C in a low calcium ACSF that contained (in mM) as follows: 130 NaCl, 11 glucose, 2.5 KCl, 2.4 MgCl_2_, 1.2 CaCl_2_, 23 NaHCO_3_, 1.2 NaH2PO_4_, and were equilibrated with 95% O2/5% CO2 and then at room temperature until the time of recording. Interleaved slices were incubated in a ACSF alone or ACSF containing 2µM URB597 for 30 minutes prior to and during the recording.

Field potential recordings were made in coronal slices containing the NAc core as previously described (7,19). Recordings were made in the medial ventral accumbens core close to the anterior commissure (7,19). For recording, slices were placed in the recording chamber and superfused (1.5 - 2 ml/min) with ACSF (same as low Ca^2+^ ACSF with the following exception: 2.4 mM CaCl_2_ and 1.2 mM MgCl_2_). All experiments were done at 25°C. Picrotoxin (100 µM) was added to the superfusion medium to block gamma-aminobutyric acid type A (GABA-A) receptors. All drugs were added at the final concentration to the superfusion medium. For field excitatory postsynaptic potential (fEPSP), the recording pipette was filled with ACSF, and afferents were stimulated with a glass electrode filled with ACSF and placed ∼200µm in the dorsal-medial direction of the recording pipette. The stimulus intensity was adjusted around 60% of maximal intensity after performing an input-output curve (baseline fEPSP amplitudes ranged between 0.15 mV and 0.4 mV). Stimulation frequency was set at 0.1 Hz. Recordings were performed with an Axopatch-200B amplifier (Axon Instrument, Molecular Device, Sunnyvale, USA). Data were lowpass filtered at 2kHz, digitized (10kHz, DigiData 1440A, Axon Instrument, Molecular Device, Sunnyvale, USA), collected using Clampex 10.2 and analyzed using Clampfit 10.2 (Axon Instrument, Molecular Device, Sunnyvale, USA). Both fEPSPs’ area and amplitude were analyzed.

### Saccharin Preference test

The test was adapted from previous studies (22) with minor modifications. Mice were individually housed and exposed to two bottles: one containing a 0.2% saccharin solution (Sigma-Aldrich, Madrid, Spain) and the other containing tap water. No previous food or water deprivation was applied before or during the test. During day 1 and 2, mice were habituated to the experimental conditions (the presence of two bottles and the saccharin taste). During the experiment, mice were injected with either cocaine or saline on day 3, and with URB597 or vehicle on day 4. Afterwards, they were placed back in their home cage. Throughout the 4-day experiment, the two bottles containing either water or saccharin solution were continuously available for the mice, except for a 30-minute period following the URB597/vehicle injection, which allowed time for URB597 to increase AEA levels (23). The consumption of water and saccharin solution was measured by weighting the bottles at 1, 2, 6 and 24 hours on days 2, 3 and 4. A control cage without animals was placed to control the amount of liquid spontaneously lost from the bottles. Saccharin preference was calculated according to the following ratio: saccharin intake(g) / [saccharin intake (g) + water intake (g)] × 100.

### Three-Chamber Social Interaction Test

The test was adapted from previous studies (24,25). Briefly, it consisted of two 10-minute phases: habituation and sociability. The box consisted of a rectangle divided into three connected chambers. The right and left chambers had a holder located in one of the corners. For habituation, mice were introduced to the central chamber and were free to move around the box. For sociability phase, a novel mouse was trapped inside one of the holders, and the experimental mouse was reintroduced into the central chamber and allowed to freely explore the box. The novel mice were male, juvenile (5-week-old), C57BL/6 mice. Time spent in each compartment and in the interactive zones (around the holders) was recorded using the SMART software (Panlab s.l.u., Barcelona, Spain) for subsequent analysis.

### qPCR

Total RNA extraction from NAc samples was conducted using the trizol method as previously described (24,26). We evaluated the expression of CB1R, FAAH, mGluR5, NAPE-PLD and GAPDH as a housekeeping gene. RT-PCR was performed by High-Capacity cDNA Reverse Transcription Kit (Applied Biosystems, Foster City, CA) using random primers and following standardized protocols. As per the qPCR, 20 ng of sample were loaded in addition to the following reagents: LightCycler SBYR green 480 Master Mix (Roche LifeScience, Product No. 04707516001) and the specific primers for the target genes (Integrated DNA Technologies, Inc.). The qPCR was performed in QuantStudio 12K Flex (Thermofisher).

### Statistical analysis

Data are presented as mean ± SEM. We used GraphPad Prism 9.5.0 and IBM SPSS Statistics 29.0 software for statistical analysis and graphing. For analysis involving two or three factors, we used two- or three-way analysis of variance (ANOVA), respectively. When an experimental condition followed a within-subject design, repeated measures ANOVA was used. The factors considered were *cocaine treatment* (cocaine/saline), *URB597 treatment* (URB597/vehicle), *time* (minutes or hours), *LTD* (baseline/LTD) and *compartment* (left/center/right, novel/center/empty, or novel/empty). The statistical threshold for significance was set at p<0.05. When F achieved significance and there was no significant variance in homogeneity, a Sidak’s *post hoc* test was run. We analyzed the results of single factor experiments with two levels using paired or unpaired two-tailed Student’s t test.

## RESULTS

### A single cocaine administration leads to loss of eCB-LTD in the NAc core

As expected, a single cocaine administration abolished plasticity in the NAc 24 hours after its administration (3) (Fig. 1B-C; Two-way ANOVA, *time* F_(49, 980)_= 51.25, p<0.001, *time* x *cocaine* F_(49, 980)_= 1.491, p=0.017), and eCB-LTD was present in the saline group (Fig. 1D; Student’s *t* test, t_(11)_= 3.426, p=0.005), but not in the cocaine group (Fig. 1E).

### A single cocaine exposure perturbates social behavior the day after use

Previous research has suggested that cocaine use alters social interaction and reward processing, but it is uncertain whether initiating cocaine use affects such behaviors in rodents beyond its acute effects. To investigate this, mice were compared in terms of their preference for consuming saccharin, a potent natural reward. Throughout the habituation phase, mice exhibited a significant inclination towards saccharin (Fig. 1F). Notably, their preference for saccharin intensified with longer periods of no alteration. Following administration of cocaine/saline, mice treated with cocaine displayed a reduction in saccharin preference (Fig 1G; Two-way ANOVA, *time* F_(3, 132)_= 92.19, p<0.001, *cocaine* F_(1, 44)_= 5.224, p=0.027, *cocaine* x *time* F_(3, 132)_= 2.687, p=0.049, Sidak 1 hour p=0.014), which was particularly noticeable at 1 hour (Fig 1H; Student’s *t* test, t_(44)_= 2.849, p=0.006), but also evident after 24 hours (Fig 1I; Student’s *t* test, t_(44)_= 2.301, p=0.026).

Next, mice were given either cocaine or saline 24 hours before being tested for social interaction. During the habituation phase, there were no group differences in preference for either compartment or locomotor activity (Fig. 1J-K). In the sociability phase, mice spent more time interacting with the novel mouse than with the empty holder (Fig. 1L; Two-way ANOVA, *compartment* F_(1, 22)_= 113.6, p<0.001) and, in particular, cocaine pre-treatment increased social interaction (Two-way ANOVA, *cocaine* F_(1, 22)_= 4.421, p=0.047).

### A single cocaine exposure decreases the NAPE-PLD gene expression levels 24 hours within administration

We investigated possible changes in the gene expression of key components of the ECS that contribute to the eCB-mediated plasticity in the NAc (Fig. 1M-P). qPCR analyses performed on the vSTR 24 hours following cocaine/saline administration revealed a reduction of NAPE-PLD gene expression in cocaine-treated mice (Fig. 1P; Student’s *t* test, t_(16)_= 2.757, p=0.013). mGluR5 gene expression levels were also reduced after cocaine administration, but the difference did not reach statistical significance (Fig. 1O; Student’s *t* test, t_(16)_= 1.819, p=0.087).

### URB597 prevents the cocaine-induced loss of eCB-LTD in the NAc core

The observed decline in NAPE-PLD and mGluR5 gene expression after the onset of cocaine use suggests an impaired mGluR5 signaling resulting in reduced AEA synthesis. Based on these findings, we hypothesized that the correction of AEA levels could reverse the synaptic and behavioral impairments caused by cocaine.

Therefore, the day after administering either cocaine or saline, we euthanized the mice and assessed the expression of eCB-LTD in the NAc in the presence of URB597. In the group that received saline, eCB-LTD was induced both in the presence or absence of URB597, although URB597 enhanced it (Fig. 2B-C; Two-way ANOVA, *time* x *URB597* F_(49, 1078)_= 1.565, p=0.008). On the other hand, in cocaine-treated mice the expression of eCB-LTD was absent, and the application of URB597 recued it (Fig. 2B and 1D; Two-way ANOVA, *URB597* F_(1, 19)_= 7.630, p=0.012, *URB597* x *time* F_(49, 931)_= 3.662, p<0.001). The comparison of the fEPSC recorded during the LTD-expression period (30-40 minutes) indicates a global effect of URB597 in enhancing LTD expression (Fig. 2E; Two-way ANOVA, *URB597* F_(1, 40)_= 9.759, p=0.003).

**Figure 2.**
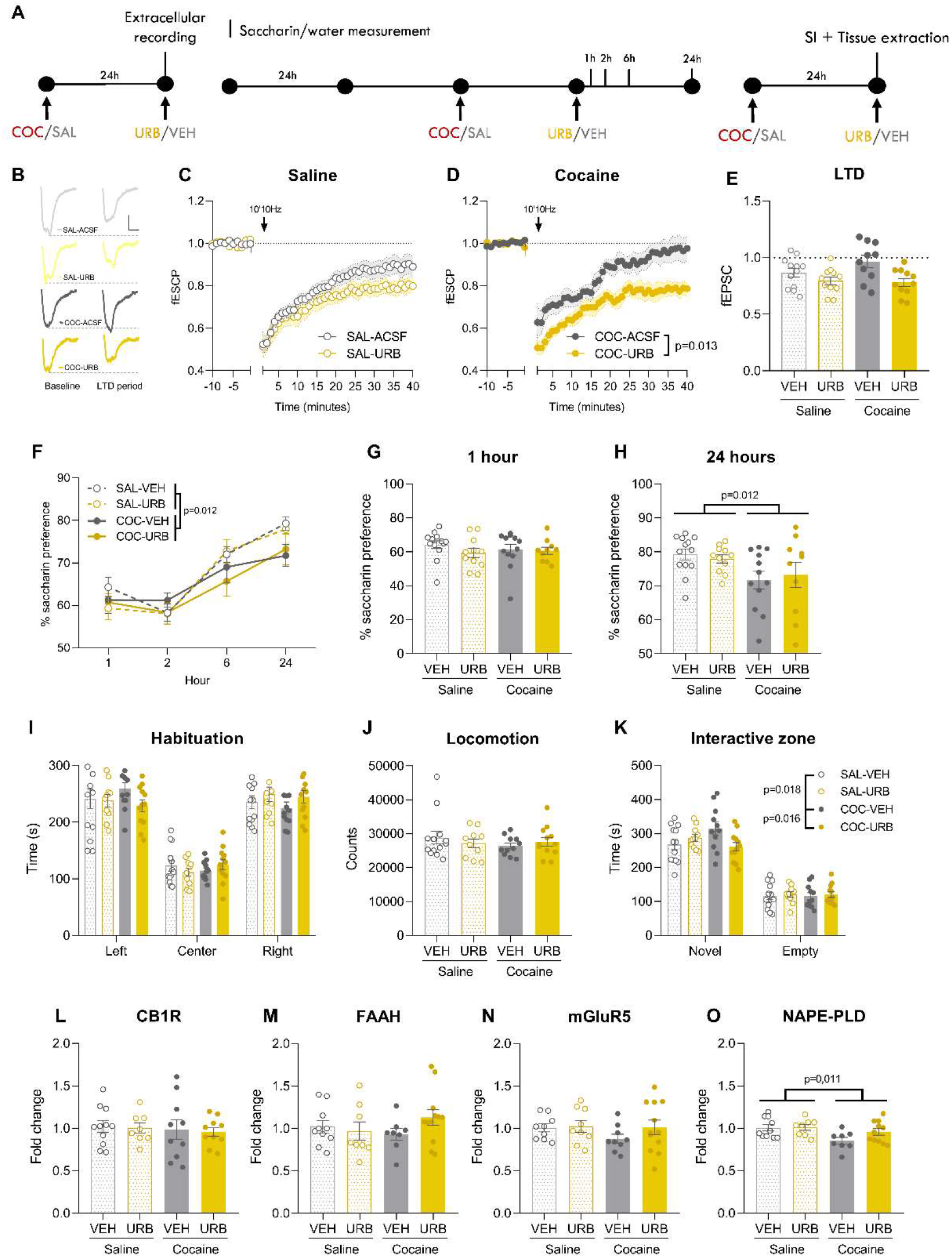
URB597 (1 mg/kg) prevents the cocaine (20 mg/kg)-induced impairment in eCB-LTD and sociability. **A** Mice were treated with cocaine/saline and 24 hours later eCB-LTD was measured with the presence of absence of URB597. Other cohorts of mice were treated with cocaine/saline and URB597/vehicle 24 hours and 30 minutes, respectively, prior to being tested saccharin preference, social behavior, and gene expression levels. **B** Representative trace during baseline (−10 to 0 minutes) and LTD expression period (30 to 40 minutes) of a SAL-treated mouse. Scale: 0.1 mV, 5 ms. **C** Average time courses of eCB-LTD for SAL-ACSF (n=13) and SAL-URB (n= 10) groups. **C** Representative trace during baseline and LTD expression period of a COC-treated mouse. Scale: 0.1 mV, 5 ms. **D** Average time courses of eCB-LTD for COC-ACSF (n=12) and COC-URB (n= 11) groups. **E** fECSPs during the LTD expression period showing that URB597 rescues the cocaine-induced loss of eCB-LTD. **F** Gene expression levels of CB1R, **G** FAAH, **H** mGluR5 and **I** NAPE-PLD 24 hours and 30 minutes following cocaine/saline and URB597/vehicle administration, respectively (SAL-VEH, n=10-11; SAL-URB, n=8-9; COC-VEH, n=8-10; COC-URB, n=10-12). **F** Cocaine-induced decrease in saccharin preference is not affected by URB597 at **G** 1 or **H** 24 hours post URB597 administration (SAL-VEH n=13, SAL-URB n=11, COC-VEH n=12 and COC-URB n=10). **I** The time spent in each compartment during the habituation phase showed that mice did not show initial preference for neither the left nor the right compartment (SAL-VEH, n=13; SAL-URB, n=10; COC-VEH, n=11; COC-URB, n=12). **J** Locomotor activity during the habituation session was not affected by either treatment. **K** Time spent in the interactive zones for each group. URB597 prevented the cocaine-induced increase in the time spent interacting with the novel mouse. **L** Gene expression levels of CB1R, **M** FAAH, **N** mGluR5 and **O** NAPE-PLD after the social interaction test (SAL-VEH, n=10-11; SAL-URB, n=8-9; COC-VEH, n=8-10; COC-URB, n=10-12). SAL: saline, COC: cocaine, VEH: vehicle, URB: URB597, SI: social interaction, LTD: long term depression, fEPSP: field postsynaptic excitatory potential, CB1R: cannabinoid receptor 1, FAAH: fatty acid amide hydrolase, mGluR5: metabotropic glutamate receptor 5, NAPE-PLD: N-acylphosphatidylethanolamine-specific phospholipase D.

### Elevation of circulating AEA levels normalizes social interaction in cocaine-exposed mice

Regarding the saccharin test, the measurements took place 24 hours following cocaine/saline administration and 30 minutes after URB597/vehicle treatment and occurred at 1, 2, 6, and 24 hours. After receiving cocaine/saline, mice treated with cocaine displayed a decrease in saccharin preference, persisting even after 48 hours (Fig. 2F-H). However, URB597 did not have any effect on saccharin preference, despite the clear effect of cocaine (Fig. 2F; Three-way ANOVA, *time* F_(3. 126)=_ 58.71, p<0.001, *cocaine* x *time* F_(3, 126)_= 3.288, p=0.023, Sidak 24 hour p=0.012; Fig. 2H; Two-way ANOVA, *cocaine* F_(1 42)_= 6.821, p=0.012).

Besides, we tested the effects of URB597 on the cocaine-induced increase in social interaction. During the habituation phase, none of the groups exhibited a preference for either the left or right compartment (Fig. 2I) or an alteration in locomotor activity (Fig. 2J). During the sociability phase, we observed that mice spent more time interacting with the novel mouse than with the empty holder (Fig. 2K; Three-way ANOVA, *compartment* F(1, 42)= 250.6, p<0.001) and, in particular, cocaine pre-treatment increased social interaction while URB597 treatment prevented such increase in the cocaine-treated mice (Three-way ANOVA, *cocaine* x *URB597* F_(2, 84)_= 312.8, p<0.001; Sidak, SAL-VEH vs COC-VEH p=0.018, COC-VEH vs COC-URB p=0.016).

### FAAH inhibition does not modify NAPE-PLD gene expression

qPCR analyses conducted on the vSTR (Fig. 2L-O) showed that cocaine administration resulted in a decrease in the gene expression of NAPE-PLD, regardless of URB597 treatment (Fig. 2O; Two-way ANOVA, *cocaine* F_(1, 33)_= 7.127, p=0.011), indicating that the URB597-induced increase in AEA is likely due to reduced degradation rather than increased synthesis.

## DISCUSSION

The current findings reveal that a single exposure to cocaine can alter saccharin preference and social behavior, as well as disrupt the neuroplastic mechanisms regulating synaptic strength in the NAc, even after the drug has been eliminated from the system. These outcomes are accompanied by a decrease in the AEA-catabolizing enzyme NAPE-PLD, suggesting a deficit in AEA signaling that could account for the suppression of plasticity induced by cocaine. Enhancing AEA levels pharmacologically through FAAH inhibition restores the cocaine-induced loss of eCB-LTD in the NAc and normalizes social behavior in cocaine-exposed mice but has no effect on the cocaine-induced decrease in saccharin preference.

Consistent with our results, cocaine-induced loss of plasticity in the NAc has been largely documented for the last 20 years, occurring after single cocaine exposure (2,3), repeated non-contingent cocaine administration (27) and cocaine self-administration (28). However, how this loss of plasticity is related behavioral alteration after the first cocaine exposure remains unclear.

We found that a single dose of cocaine leads to a reduction in saccharin intake for the subsequent 48 hours. While the effects observed during the first hour after cocaine administration can be attributed to the stimulant’s impact on locomotor activity, which may interfere with consumption behaviors, the extended suppressive effects beyond this time frame cannot be explained by this mechanism. Instead, we hypothesize that cocaine exposure induces long-lasting neuroplastic adaptations in the mesocorticolimbic system, responsible for natural reward processing, leading to the observed changes in saccharin preference. In line with this, previous studies have extensively documented that pairing cocaine with saccharin dramatically decreases saccharin intake (12,13,29–31). This effect appears to be due to a decrease in the perceived value of saccharin as a reward, rather than an impairment in sweet taste sensitivity (13). The ability of cocaine to decrease natural reward is supported by studies using intracranial self-stimulation. While cocaine appears to facilitate brain reward function when present in the (32,33), the opposite effect is observed during withdrawal (33,34). Interestingly, pharmacological (35) or genetic deletion of mGluR5 in D1-expressing SPNs (11) also increases intracranial self-stimulation thresholds, suggesting impaired reward function. This evidence supports the idea that cocaine decreases saccharin preference by disrupting natural reward function, likely due in part to an impairment in mGluR5 signaling. Alternatively, cocaine has been observed to have a negative impact on appetite (36,37), often resulting in significant weight loss among cocaine users (38). This could potentially contribute to the decrease in saccharine preference caused by cocaine. However, it is still uncertain whether this effect could account for the continued decrease in saccharine preference observed up to 48 hours after cocaine administration.

Furthermore, a single dose of cocaine led to increased social interaction 24 hours after administration. Given that the half-life of cocaine in the mouse brain is 16 minutes (39), this behavioral finding cannot be attributed to the drug’s acute effects. Instead, it implies that the central nervous system undergoes neuroplastic adaptations due to cocaine administration, leading to behavioral changes even when the drug is no longer present. In contrast, previous studies have indicated that social interaction is reduced during acute (14–16) or repeated non-contingent cocaine administration (17). These discrepancies suggest that the immediate behavioral effects of the drug are distinct from those produced by cocaine-induced neuroplastic changes. Wang et al. (2014) reported decreased social interaction after a 4-day cocaine treatment (20 mg/kg) when measured 24 hours after the last administration (18), contrasting with our findings. Possible explanations for these disparities include the use of different measurement models for social behaviors (three-chamber test versus open arena) or varying effects of cocaine on social behavior depending on whether it is administered acutely or repeatedly.

Cocaine-treated mice showed a decrease in NAPE-PDL gene expression levels after undergoing the social interaction test. Similarly, acute cocaine administration was associated with a decrease in gene and protein expression of NAPE-PDL and DAGLα in the hippocampus, resulting in an overall reduction in eCB synthesis (40). Based on these findings, we hypothesize that a single exposure to cocaine may result in a reduction in the production of AEA, thus playing a role in the impairment of eCB-LTD in the NAc. Therefore, we pharmacologically inhibited FAAH activity with URB597 to test whether an increase in AEA levels could restore the cocaine-induced plasticity impairments as well as behavioral alterations.

We found that the enhancement of AEA caused by FAAH inhibition is sufficient to restore the loss of eCB-LTD after a single cocaine exposure, indicating a prominent role of AEA in mediating this kind of plasticity. Congruently, Grueter et al (2010) found an enhancement of both TRPV1- and CB1R-mediated LTD in the NAc following URB597 administration (2). Considering that the function of TRPV1 and CB1R appears unaffected following cocaine exposure (2,41) and that mGluR5 signaling is hindered due to receptor internalization by the homer scaffolding proteins (3,42), it is proposed that the decline in eCB-LTD observed after cocaine use is primarily due to a reduction in mGluR5’s capability to convert anterograde glutamate transmission into eCB production, including AEA. However, a complementary role of 2-AG in mediating this type of plasticity and the cocaine-induced alterations cannot be ruled out. Based on these observations, we propose that a single cocaine exposure disrupts eCB-LTD by impairing mGluR5 signaling and decreasing AEA synthesis. Reduced AEA levels lead to decreased activation of both presynaptic CB1R, resulting in a blunted inhibition of glutamate release, and postsynaptic TRPV1, causing an impaired AMPAR internalization (Fig. 3). It is yet to be determined whether the effects of AEA are primarily targeted towards CB1R or TRPV1, or whether other NAEs also play a role in this process.

**Figure 3.**
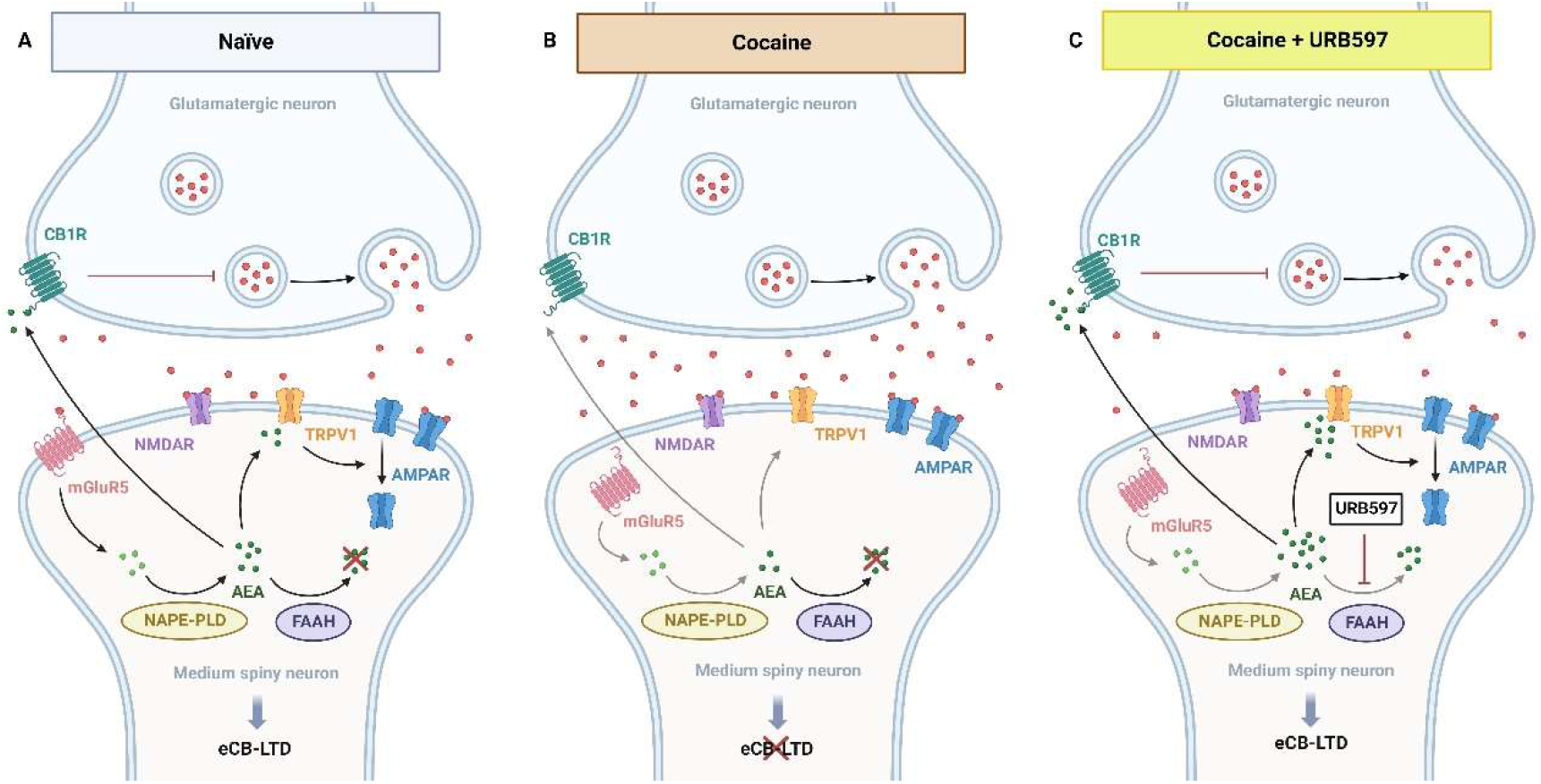
URB597 actions in the NAc synapses. **A** In naïve mice, eCB-LTD exerts negative control over glutamatergic synapses in the NAc through mGluR5 signaling. When glutamate binds to mGluR5, it triggers the production of AEA, which in turn suppresses glutamate release through CB1R binding and causes AMPAR internalization through TRPV1 signaling. **B** One day following cocaine exposure, mGluR5 internalization impairs the downstream pathway and suppresses AEA release, resulting in a loss inhibitory control over SPNs activation. **C** The inhibition of AEA degradation by URB597 rescues the AEA-dependent activation of CB1R and TRPV1R and restores eCB-LTD in the NAc. eCB-LTD: endocannabinoid-mediated long term depression, CB1R: cannabinoid receptor 1, NMDAR: N-methyl-D-aspartate receptor, AMPAR: α-amino-3-hydroxy-5-methyl-4-isoxazolepropionic acid receptor, TRPV1: transient receptor potential vanilloid 1, mGluR5: metabotropic glutamate receptor 5, FAAH: fatty acid amide hydrolase, NAPE-PLD: N-acylphosphatidylethanolamine-specific phospholipase D, AEA: anandamide. Created with biorender.com.

However, URB597 treatment did not affect cocaine-induced suppression of saccharin intake. A previous study reported that URB597 (0.5 mg/kg) reduced saccharin preference in mice (43), which is consistent with the finding that URB597 raises intracranial self-stimulation thresholds (44). Such effect was not observed here as URB597 did not affect saccharin preference in either cocaine- or saline-treated mice. The reasons why an increase in AEA appears to prevent cocaine-induced changes in social interaction but not saccharin preference remain unclear. One possible hypothesis is that the brain processes these two types of natural rewards differently, and therefore, they rely on distinct neural circuits that may involve varying levels of AEA in the synapses. Thus, AEA may play a different role in the neural pathways involved in processing social rewards versus those involved in processing sweet rewards.

In the case of social behavior, we found that URB597 treatment 30 minutes before the test prevented the cocaine-induced increase in sociability. To our knowledge, this is the first evidence of the impact of FAAH inhibition on social interaction after cocaine administration. Prior examinations of the role of URB597 on sociability have yielded inconclusive results. While some studies reported a URB597-induced decline in social behavior (45–47), others found the opposite outcome (48,49). In line with the present results, two independent studies reported a reduction in social interaction following systemic administration of URB597 that is mediated through TRPV1 rather than CB1R (46,47). Moreover, an effective reduction in operant responding for social play was observed with a dosage of 0.2 mg/kg of URB597 (50). Further research is required to elucidate the involvement of AEA in cocaine-induced alterations in social behavior, as well as the precise mechanisms underlying such modulation.

The present study has two main limitations. Firstly, while the impairments induced by cocaine in both accumbal plasticity and related behaviors follow the same administration regimen, the current experimental design does not allow us to determine a causal relationship between these outcomes. Therefore, it is crucial to experimentally manipulate the expression of eCB-LTD(11) after cocaine exposure in order to uncover whether the observed behavioral alterations are dependent on this plasticity. Secondly, conducting a broader pharmacological examination by using antagonists targeting different NAEs receptors, such as CB1R or TRPV1, would provide valuable information regarding the mechanism through which URB597 restores eCB-LTD.

In conclusion, the results of the present study indicate that acute cocaine administration enhances social behavior and reduces saccharin preference after the drug has left the system. Parallelly, a single dose of cocaine disrupts the neuroplastic mechanisms that control synaptic strength in the NAc and decreases the gene expression of the AEA-catabolizing enzyme NAPE-PLD. We evidenced that the cocaine-induced loss of eCB-LTD is mediated by AEA, as FAAH inhibition is sufficient to restore the impairment caused by cocaine. Furthermore, FAAH inhibition normalizes the cocaine-induced alterations in social behavior. Future investigations should establish whether the effects of cocaine and URB597 on eCB-LTD and social behavior are causally connected. Overall, this study demonstrates that a single exposure to cocaine can induce neuroplastic changes that result in behavioral alterations even in the absence of the drug and highlights the involvement of the ECS, especially AEA, in the early impairments induced by cocaine.

## Acknowledgements

We thank Inés Gallego-Landín, Mireia Medrano and Xavier Puig-Reyné for his assistance carrying out the experiment and Adriana Castro-Zavala for her assistance in the use of BioRender.

This work was supported by the Ministerio de Ciencia e Innovación (#PID2019-104077RB-100 MCIN/ AEI/10.13039/501100011033), Ministerio de Sanidad (Plan Nacional sobre Drogas #2018/007 and ISCIII-Feder-RIAPAd-RICORS #RD21/0009/001), the Generalitat de Catalunya, AGAUR (#2021SGR00485) and by the Institut National de la Santé et de la Recherche Médicale (INSERM). L.A-Z received a FPI grant (BES-2017-080066) associated to #SAF2016-75966-R grant to O.V. from Ministerio de Economia y Competitividad. P.B-S received a FI-AGAUR grant from the Generalitat de Catalunya (2021FI_B00205). The Department of Medicine and Health Sciences (UPF) is a “Unidad de Excelencia María de Maeztu” funded by the AEI (#CEX2018-000792-M). O.V. is recipient of an ICREA Academia Award (Institució Catalana de Recerca i Estudis Avançats, Generalitat de Catalunya).

## REFERENCES

1. Lüscher C, Robbins TW, Everitt BJ (2020): The transition to compulsion in addiction. Nature Rev Neurosci 21: 247–263.

2. Grueter BA, Brasnjo G, Malenka RC (2010): Postsynaptic TRPV1 triggers cell type–specific long-term depression in the nucleus accumbens. Nat Neurosci 13: 1519–1525.

3. Fourgeaud L, Mato S, Bouchet D, Hémar A, Worley PF, Manzoni OJ (2004): A single In Vivo exposure to cocaine abolishes endocannabinoid-mediated long-term depression in the nucleus accumbens. J Neurosci 24: 6939–6945.

4. Parsons LH, Hurd YL (2015): Endocannabinoid signalling in reward and addiction. Nature 16: 579–594.

5. Grueter BA, Rothwell PE, Malenka RC, Sheng M, Triller A (2012): Integrating synaptic plasticity and striatal circuit function in addiction. Curr Opin Neurobiol 22: 545–551.

6. Cristino L, Bisogno T, Di Marzo V (2019): Cannabinoids and the expanded endocannabinoid system in neurological disorders. Nat Rev Neurol 16: 9–29.

7. Robbe D, Kopf M, Remaury A, Bockaert J, Manzoni OJ (2002): Endogenous cannabinoids mediate long-term synaptic depression in the nucleus accumbens. Proc Natl Acad Sci U S A 99: 8384–8388.

8. Castillo PE, Younts TJ, Chávez AE, Hashimotodani Y (2012): Endocannabinoid signaling and synaptic function. Neuron 76: 70.

9. Van Der Stelt M, Marzo V Di (2005): Anandamide as an intracellular messenger regulating ion channel activity. Prostaglandins Other Lipid Mediat 77: 111–122.

10. Smart D, Gunthorpe MJ, Jerman JC, Nasir S, Gray J, Muir AI, et al. (2000): The endogenous lipid anandamide is a full agonist at the human vanilloid receptor (hVR1). Br J Pharmacol 129: 227–230.

11. Bilbao A, Neuhofer D, Sepers M, Wei S peng, Eisenhardt M, Hertle S, et al. (2020): Endocannabinoid LTD in Accumbal D1 Neurons Mediates Reward-Seeking Behavior. iScience 23: 100951.

12. Freet CS, Steffen C, Nestler EJ, Grigson PS (2009): Overexpression of ΔFosB Is Associated With Attenuated Cocaine-Induced Suppression of Saccharin Intake in Mice. Behav Neurosci 123: 397–407.

13. Roebber JK, Izenwasser S, Chaudhari N (2015): Cocaine decreases saccharin preference without altering sweet taste sensitivity. Pharmacol Biochem Behav 133: 18–24.

14. Estelles J, Rodríguez-Arias M, Aguilar MA, Miñarro J (2004): Social behavioural profile of cocaine in isolated and grouped male mice. Drug Alcohol Depend 76: 115–123.

15. Rademacher DJ, Schuyler AL, Kruschel CK, Steinpreis RE (2022): Effects of cocaine and putative atypical antipsychotics on rat social behavior An ethopharmacological study. Pharmacol Biochem Behav 73: 769–778.

16. Lluch J, Rodríguez-Arias M, Aguilar MA, Miñarro J (2005): Role of dopamine and glutamate receptors in cocaine-induced social effects in isolated and grouped male OF1 mice. Pharmacol Biochem Behav 82: 478–487.

17. Morisot N, Monier R, Le Moine C, Millan MJ, Contarino A (2018): Corticotropin-releasing factor receptor 2-deficiency eliminates social behaviour deficits and vulnerability induced by cocaine. Br J Pharmacol 175: 1504–1518.

18. Wang J-L, Wang B, Chen W (2014): Differences in cocaine-induced place preference persistence, locomotion and social behaviors between C57BL/6J and BALB/cJ mice. Zool Res 35: 426–435.

19. Neuhofer D, Lassalle O, Manzoni OJ (2018): Muscarinic M1 Receptor Modulation of Synaptic Plasticity in Nucleus Accumbens of Wild-Type and Fragile X Mice. ACS Chem Neurosci 9: 2233–2240.

20. Jung KM, Sepers M, Henstridge CM, Lassalle O, Neuhofer D, Martin H, et al. (2012): Uncoupling of the endocannabinoid signalling complex in a mouse model of fragile X syndrome. Nat Commun 3: 1080.

21. Lafourcade M, Larrieu T, Mato S, Duffaud A, Sepers M, Matias I, et al. (2011): Nutritional omega-3 deficiency abolishes endocannabinoid-mediated neuronal functions. Nat Neurosci 14: 345–350.

22. Gracia-Rubio I, Moscoso-Castro M, Pozo OJ, Marcos J, Nadal R, Valverde O (2016): Maternal separation induces neuroinflammation and long-lasting emotional alterations in mice. Prog Neuropsychopharmacol Biol Psychiatry 65: 104–117.

23. Fegley D, Gaetani S, Duranti A, Tontini A, Mor M, Tarzia G, Piomelli D (2005): Characterization of the Fatty Acid Amide Hydrolase Inhibitor Cyclohexyl Carbamic Acid 3-Carbamoyl-biphenyl-3-yl Ester (URB597): Effects on Anandamide and Oleoylethanolamide Deactivation. J Pharmacol Exp Ther 313: 352–358.

24. Castro-Zavala A, Alegre-Zurano L, Cantacorps L, Gallego-Landin I, Welz PS, Benitah SA, Valverde O (2022): Bmal1-knockout mice exhibit reduced cocaine-seeking behaviour and cognitive impairments. Biomed Pharmacother 153: 113333.

25. Alegre-Zurano L, Martín-Sánchez A, Valverde O (2020): Behavioural and molecular effects of cannabidiolic acid in mice. Life Sci 259: 118271.

26. Castro-Zavala A, Martín-Sánchez A, Montalvo-Martínez L, Camacho-Morales A, Valverde O (2021): Cocaine-seeking behaviour is differentially expressed in male and female mice exposed to maternal separation and is associated with alterations in AMPA receptors subunits in the medial prefrontal cortex. Prog Neuropsychopharmacol Biol Psychiatry 109: 110262.

27. Huang C-C, Yeh C-M, Wu M-Y, Chang AYW, Chan JYH, Chan SHH, Hsu K-S (2011): Cocaine Withdrawal Impairs Metabotropic Glutamate Receptor-Dependent Long-Term Depression in the Nucleus Accumbens. J Neurosci 31: 4194–4203.

28. Martin M, Chen BT, Hopf FW, Bowers MS, Bonci A (2006): Cocaine self-administration selectively abolishes LTD in the core of the nucleus accumbens. Nat Neurosci 9: 868–869.

29. Freet CS, Lawrence AL (2015): Ceftriaxone attenuates acquisition and facilitates extinction of cocaine-induced suppression of saccharin intake in C57BL/6J mice. Physiol Behav 149: 174–180.

30. Twining RC, Freet CS, Wheeler RA, Reich CG, Tompers DA, Wolpert SE, Grigson PS (2016): The role of dose and restriction state on morphine-, cocaine-, and LiCl-induced suppression of saccharin intake: A comprehensive analysis. Physiol Behav 161: 104–115.

31. Grigson PS, Twining RC (2002): Cocaine-induced suppression of saccharin intake: A model of drug-induced devaluation of natural rewards. Behav Neurosci 116: 321–333.

32. Fish EW, Riday TT, Mcguigan MM, Faccidomo S, Hodge CW, Malanga CJ (2010): Alcohol, Cocaine, and Brain Stimulation-Reward in C57Bl6/J and DBA2/J Mice. Alcohol Clin Exp Res 34: 81–89.

33. Stoker AK, Markou A (2011): Withdrawal from Chronic Cocaine Administration Induces Deficits in Brain Reward Function in C57BL/6J Mice. Behav Brain Res 223: 176–181.

34. Stoker AK, Olivier B, Markou A (2012): Involvement of metabotropic glutamate receptor 5 in brain reward deficits associated with cocaine and nicotine withdrawal and somatic signs of nicotine withdrawal. Psychopharmacology (Berl) 221: 317–327.

35. Cleva RM, Watterson LR, Johnson MA, Olive MF (2012): Differential modulation of thresholds for intracranial self-stimulation by mGlu5 positive and negative allosteric modulators: Implications for effects on drug self-administration. Front Pharmacol 2: 93.

36. Murphy KG (2005): Dissecting the role of cocaine- and amphetamine-regulated transcript (CART) in the control of appetite. Brief Funct Genomic Proteomic 4: 95–111.

37. Billing L, Ersche KD (2015): Cocaine’s appetite for fat and the consequences on body weight. Am J Drug Alcohol Abuse 41: 115–118.

38. Ersche KD, Stochl J, Woodward JM, Fletcher PC (2013): The skinny on cocaine: insights into eating behavior and body weight in cocaine-dependent men. Appetite 71: 75–80.

39. Benuck M, Lajtha A, Reith MEA (1987): Pharmacokinetics of Systemically Administered Cocaine and Locomotor Stimulation in Mice. J Pharmacol Exp Ther 243: 144–149.

40. Blanco E, Galeano P, Palomino A, Pavón FJ, Rivera P, Serrano A, et al. (2016): Cocaine-induced behavioral sensitization decreases the expression of endocannabinoid signaling-related proteins in the mouse hippocampus. Eur Neuropsychopharmacol 26: 477–492.

41. Mccutcheon JE, Loweth JA, Ford KA, Marinelli M, Wolf ME, Tseng KY (2011): Group I mGluR Activation Reverses Cocaine-Induced Accumulation of Calcium-Permeable AMPA Receptors in Nucleus Accumbens Synapses via a Protein Kinase C-Dependent Mechanism. J Neurosci 31: 14536–14541.

42. Hoffmann HM, Crouzin N, Moreno E, Raivio N, Fuentes S, McCormick PJ, et al. (2017): Long-lasting impairment of mGluR5-activated intracellular pathways in the striatum after withdrawal of cocaine self-administration. Int J Neuropsychopharmacol 20: 72–82.

43. Zhou Y, Schwartz BI, Giza J, Gross SS, Lee FS, Kreek MJ (2017): Blockade of alcohol escalation and “relapse” drinking by pharmacological FAAH inhibition in male and female C57BL/6J mice. Psychopharmacology (Berl) 234: 2955–2970.

44. Vlachou S, Nomikos GG, Panagis G (2006): Effects of endocannabinoid neurotransmission modulators on brain stimulation reward. Psychopharmacology (Berl) 188: 293–305.

45. Matricon J, Seillier A, Giuffrida A (2016): Distinct neuronal activation patterns are associated with PCP-induced social withdrawal and its reversal by the endocannabinoid-enhancing drug URB597. Neurosci Res 110: 49–58.

46. Seillier A, Giuffrida A (2018): The cannabinoid transporter inhibitor OMDM-2 reduces social interaction: Further evidence for transporter-mediated endocannabinoid release. Neuropharmacology 130: 1–9.

47. Seillier A, Martinez AA, Giuffrida A (2013): Phencyclidine-Induced Social Withdrawal Results from Deficient Stimulation of Cannabinoid CB 1 Receptors: Implications for Schizophrenia. Neuropsychopharmacology 38: 1816–1824.

48. Manduca A, Servadio M, Campolongo P, Palmery M, Trabace L, JMJ Vanderschuren L, et al. (2014): Strain-and context-dependent effects of the anandamide hydrolysis inhibitor URB597 on social behavior in rats. Eur Neuropsychopharmacol 24: 1337–1348.

49. Trezza V, Damsteegt R, Manduca A, Petrosino S, Van Kerkhof LWM, Pasterkamp RJ, et al. (2012): Endocannabinoids in Amygdala and Nucleus Accumbens Mediate Social Play Reward in Adolescent Rats. J Neurosci 32: 14899–14908.

50. Marijke Achterberg E, van Swieten MM, Driel N V, Trezza V, JMJ Vanderschuren L (2016): Dissociating the role of endocannabinoids in the pleasurable and motivational properties of social play behaviour in rats. Pharmacol Res 110: 151–158.

